# Abundant and active acetogens enhance the carbon dioxide sink of Blue Carbon ecosystems

**DOI:** 10.1101/2025.01.07.631696

**Authors:** Karen Rodriguez, Francesco Ricci, Gaofeng Ni, Naima Iram, Robin Palfreyman, Ricardo A. Gonzalez-Garcia, James Heffernan, Chris Greening, Maria Fernanda Adame, Esteban Marcellin

## Abstract

Blue Carbon ecosystems, which include all tidal wetlands, mitigate climate change by capturing and storing carbon dioxide (CO_2_) from the atmosphere. Most carbon fixation in these systems is thought to be driven by plant and microbial photosynthesis, whereas chemosynthetic processes are assumed to play a minor role. However, these ecosystems often contain anoxic environments ideal for chemosynthetic microbes such as acetogens. Here, we show that acetogens are abundant and active mediators of carbon sequestration in tidal wetland soils by pairing gene-and genome-resolved metagenomic analysis with isolation and analysis of gas-fermenting acetogens in bioreactors. Metagenomic profiling revealed that diverse microbes can mediate carbon fixation, primarily through the Calvin-Benson-Bassham cycle and Wood-Ljungdahl pathways. These include various bacteria and archaea capable of reductive acetogenesis. On this basis, we grew bacterial enrichment cultures from tidal wetland soils using the gases hydrogen and CO_2_ as the sole energy and carbon sources. Bioreactor analysis revealed that these enrichments are dominated by clostridial acetogens that grow rapidly by converting CO_2_ into acetate and other products. Collectively, these results reveal Blue Carbon ecosystems harbour communities that can exclusively subsist by using CO_2_ as their sole electron acceptor and for carbon fixation, thereby providing evidence of a novel carbon sink pathway within these ecosystems beyond the known mechanisms of photosynthetic carbon fixation and soil sequestration. Additionally, the discovery and isolation of these chemosynthetic communities provide opportunities for developing further mechanisms of CO_2_ removal through industrial gas fermentation.

## Introduction

Addressing climate change represents an urgent challenge for humanity. This endeavour requires the exploration and implementation of nature-based solutions that act synergistically with efforts to mitigate fossil fuel emissions. Blue Carbon ecosystems sequester carbon dioxide (CO_2_), thus offering significant greenhouse gas removals that align with climate change adaptation and mitigation policies [1, 2]. Despite coastal and marine blue carbon ecosystems occupying less than 2% of the global ocean area, they account for ∼50% of carbon storage in its sediments [3]. Beyond carbon sequestration, these ecosystems hold significant ecological and economic value, providing crucial services such as habitat for commercial fisheries, enhanced water quality, and protected coastlines [4]. The carbon sink capability of Blue Carbon ecosystems is driven by a complex interplay of biological, chemical, and physical factors. Among these ecosystems, mangroves have the highest carbon stocks [5], characterised by elevated soil respiration concomitant with dynamic CO_2_ and methane (CH_4_) fluxes. Tidal salt marshes, where periodic to continuous flooding slows decomposition processes, can also be highly productive and exceed the carbon sink potential of many terrestrial forests [6–10].

Various metabolic pathways and organisms contribute to CO_2_ sequestration in Blue Carbon ecosystems. The main CO_2_ fixation mechanism is photosynthesis, which drives the accumulation of organic carbon as woody biomass (trunks, branches, large roots) and organic material (leaves, detritus, fine roots) sequestered in the soils [11]. However, some tidal wetlands (salt marshes and forests dominated by *Melaleuca* spp.) exhibit CO_2_ uptake in dark conditions [12, 13], suggesting the presence of carbon fixation pathways independent of light. Recent metagenomic studies have suggested chemosynthetic microorganisms enhance CO_2_ fixation in estuarine and coastal wetland ecosystems, indicating the potential presence and chemolithoautotrophic activity of members from the phyla Desulfobacterota, Gemmatimonadota, Methylomirabilota, Nitrospirota and Proteobacteria within these ecosystems [14–16]. It has been estimated that chemolithoautotrophs in estuarine and coastal ecosystems sequester 328 Tg C year^-1^, accounting for ∼42.6% of total oceanic dark carbon fixation [17]. In wetlands, the metabolism of chemoautotrophs is facilitated by pronounced vertical physicochemical gradients (e.g., salinity, ammonium, pH, oxygen) [18, 19].

These gradients drive the spatial segregation of microbial functional guilds [20, 21], with oxygen sensitive metabolisms such as fermentation, sulfate reduction, and methanogenesis generally thought to be confined to deeper anaerobic niches [22]. The waste products of these metabolisms provide niches for chemolithoautotrophs, which utilise inorganic and one carbon compounds such as H_2_, H_2_S and CH_4_ for energy conservation and growth. Together, this interplay between physicochemical and biological factors ultimately produces an environment where CO_2_ is efficiently sequestered at the soil interface, enhancing the greenhouse gas mitigating capacity of these environments.

Acetogens are an understudied yet crucial group of carbon-fixing microorganisms in Blue Carbon ecosystems [14, 16, 23]. These microbes are particularly notable for their ability to convert CO_2_ into acetate through the Wood-Ljungdahl pathway, considered the most efficient carbon fixation pathway [24, 25]. Under anoxic conditions, this pathway results in the reduction of two molecules of CO_2_ to acetyl-CoA, coupled to the oxidation of electron donors such as H_2_ into acetyl-CoA. In addition to being highly efficient, acetogens are also highly metabolically versatile, enabling them to convert diverse substrates available into acetate [26]. Their activity contributes to microbial community dynamics, often engaging in cross-feeding interactions. For example, acetogens may consume H_2_ produced by fermenters or supply acetate as substrate to methanogens, highlighting their central roles in microbial interactions and food webs across various environments [27, 28]. Previous studies have shown that the marker genes associated with acetogenesis and other chemosynthetic processes are present in Blue Carbon ecosystems [14, 15, 29, 30]. This suggests acetogens potentially contribute to the CO_2_ sequestration capacity of tidal wetlands, especially at subsurface sediments, where light-dependent photosynthesis and oxygen levels are limited. However, genome-and activity-based evidence supporting their contribution to dark carbon fixation within wetland ecosystems is lacking, underscoring the need for their characterisation and further investigation of their carbon fixation capabilities and ecological role.

In this study, we investigated the presence of CO_2_-fixing bacterial communities in tidal salt marshes and *Melaleuca* forests from a subtropical tidal wetland in Australia. Based on negative dark CO_2_ fluxes observed in these ecosystems [12], we hypothesized that microbes could play an important role through chemosynthetic CO_2_ fixation. To test our hypothesis, we first profiled the composition and capabilities of the CO_2_-fixing microorganisms in these environments through gene-and genome-resolved metagenomics, then isolated and characterised microbes capable of growing on C1 gases and H_2_ as only carbon and energy sources. Through these approaches, we infer acetogens are abundant, widespread, and active in tidal wetlands and likely other Blue Carbon ecosystems. In turn, these findings emphasize that diverse and uncultured microorganisms play a key role in mediating carbon fixation within Blue Carbon ecosystems. This underscores the importance of considering such microorganisms when quantifying and managing oceanic carbon sinks and highlights their potential applications in gas fermentation technologies.

## Materials and Methods

### Site description and determination of CO_2_ sequestration in wetlands

The samples analysed in this study were taken from the Yandina Wetlands Restoration Project within the Maroochy River catchment, Queensland, Australia. This area has a humid subtropical climate, with a mean annual rainfall of 1,706 mm (highest mean rainfall of 257 mm in February and lowest of 54 mm in August; Station 40157, 1896-2015, Australian Bureau of Meteorology, ABM, 2021). The area has two main seasons: a dry, cool season (April to September: 26.0 to 23.1°C) and a humid, warm season (October to March: 25.6 to 27.9 °C, ABM, 2021). The site covers 191 ha from the town of Coolum, bordered by Small Creek on the southern side and Yandina Creek on the northern side. The Yandina Wetlands Restoration Project is part of the “Blue Heart initiative” that aims to restore and protect wetlands in an area of 5,000 ha, mainly to manage flooding and improve water quality, enhance carbon sequestration, increase biodiversity, and provide recreational opportunities. The site is a mosaic of tidal wetland vegetation composed of supratidal forests, including *Melaleuca* and *Casuarina spp* forests, salt marshes and mangroves. Greenhouse gas fluxes were obtained from Iram et al. 2021, where CH_4_ and CO_2_ soil fluxes were measured every three months between July 2018 and July 2019, using closed chambers made of round polyvinyl chloride (PVC) pipes [31]. Because the chambers were dark, we only accounted for CO_2_ soil respiration and uptake, excluding photosynthesis.

### Sample collection and metagenomic sequencing

Soils samples of the Yandina tidal wetlands were collected in July 2019 from two locations, respectively covered by salt marsh (26°33.800’S, 153°02.663’E) and emerging *Melaleuca* woodlands (26°33.708’S, 153°02.064’E), at depths of 5 and 7.5 cm. Samples were transported to our laboratories under anoxic conditions and stored at -80 °C until further processing. Genomic DNA for microbial community identification was recovered from 250 mg of wet soil from each sample using the DNeasy PowerSoil Pro Kit (Qiagen, Germany). DNA quantity and purity was measured using a NanoDrop-2000 spectrophotometer (Thermo Fisher, USA). Recovered DNA was subject to metagenomic sequencing by the Ramaciotti Centre for Genomics on a NextSeq 1000 (Illumina, USA) platform to produce 2x150 bp paired-end reads.

### Metabolic annotation of metagenomic short reads

Contaminating PhiX and low-quality sequences (minimum quality score 20) were removed from the four metagenome samples generated in this study plus a total of 27 publicly available metagenomes from various Blue Carbon ecosystems including seagrass [32, 33], saltmarsh [34] and mangroves [35] using the BBDuk function of BBTools v.36.92 (https://sourceforge.net/projects/bbmap/). The quality-filtered forward reads with lengths of at least 100 bp were then examined for the presence of the marker genes using DIAMOND v.0.9.31 [36]. Specifically, reads were compared against the custom-made reference databases of 51 metabolic marker genes [37] for energy conservation, carbon fixation, phototrophy, and hydrogen, carbon monoxide, methane, sulfur, nitrogen, and iron cycling. A query coverage of 80% and an identity threshold of 80% (for *psaA*), 75% (for *hbsT*), 70% (for *atpA, psbA, isoA, ygfK, aro*), 60% (for *amoA, mmoA, coxL, FeFe, nxrA, rbcL, nuoF*) and 50% (for all others) was used. Read counts were normalized to reads per kilobase per million (RPKM). To infer abundance, read counts were further normalized to gene length and the abundance of single-copy marker genes.

### Microbial isolation

Microbes were isolated using culture dilution techniques with gaseous substrate selective pressures, followed by spread-plate methods. Soil samples were first homogenized in chemically defined PETC-MES media supplemented with 1.5 g/L yeast extract [38]. The homogenous mixtures (2.5 mL) were inoculated into an anaerobic serum bottle containing 25 mL of fresh liquid media and incubated in a shaking incubator at 150 rpm, 37°C, and pH 5.7. Cultures were supplemented with 2 mM 2-bromoethanesulfonate (BES) as a methanogen inhibitor and gas-pressurized at 190 kPa. Three gas mixtures were employed: 2% CO / 23% CO_2_ / 65% H_2_ (syngas), 60% CO, and 23% CO_2_ / 67% H_2_, with all gas mixes balanced to 100% with Argon. Bacterial growth and gas consumption over time were analysed. Cultures grown on CO_2_/H_2_ were consecutively sub-cultured into fresh gas and media four times, as these were the only cultures with decreased pressure in the headspace. Agar plates were prepared with PETC-MES media supplemented with 2 g/L sodium bicarbonate, 1.5 g/L yeast extract, 2.1 g/L BES, 10 g/L agar. The 2nd, 3rd, and 4th sub-cultures (200 µl each) were streaked onto agar plates and incubated for five weeks at 37°C in an anaerobic chamber pressurized with 70 kPa of the CO_2_/H_2_ gas mixture. Four colonies from each plate were selected based on morphological differences and cultured anaerobically in liquid media using the abovementioned conditions. Two cultures, named A and B, demonstrating increased growth and gas consumption, were sub-cultured into fresh media and gas, and stored in 25% glycerol cryovials at -80°C. Samples for metagenomics sequencing were obtained from enriched bioreactor cultures A and B. Genomic DNA extraction, metagenomic library preparation and paired-end sequencing was performed by the Australian Centre for Ecogenomics using a NovaSeq 600 (Illumina, USA) platform.

### Bioreactor batch fermentation of enrichment cultures

The growth, extracellular product profile, and gas consumption of cultures A and B were analysed in batch bioreactors for 171 h. Before bioreactor inoculation, pre-cultures were grown in serum bottles using the abovementioned conditions and sub-cultured twice. The cultures were then inoculated at an initial OD of 0.1 into 500 mL reactor vessels (Infors HT, Switzerland) with modified PETC-MES media supplemented with 1.5 g/L of yeast extract [39]. Fermentation conditions included: continuous gas flow of the CO_2_/H_2_ mixture at 8 mL/min (∼1 atm in headspace), controlled by a mass flow controller, pH 5.5 automatically maintained using 1 M NaOH and temperature controlled at 37°C. Stirring was increased twice at 120 and 143 h to improve gas transfer into the liquid culture.

### Analytical methods for growth, substrate and product measurements

Biomass analyses of all cultures were conducted through OD_600_ measurements using a Libra S12 UV/Vis spectrophotometer (Biochrom, UK). Specific growth rates (µ_max_) were calculated using three data points from the exponential phase, yielding a correlation coefficient R^2^ > 0.99 between culture time and natural logarithm of OD_600_. The pressure of CO_2_/H_2_ gases in the headspace of serum bottle cultures was measured using a manometer (Omega, USA) and sterile needle connection. In bioreactor cultures A and B, off-gases were quantified at high resolution with an online HPR-20-QIC mass spectrometer (Hiden Analytical, UK). The intensities of H_2_, Ar, and CO_2_ were monitored by a Faraday cup at 2, 40 and 44 amu. These masses were chosen so that a unique signal would represent each target compound. To achieve reliable off-gas measurements, the cylinder was used as calibration gas in each analysis cycle from the mass spectrometer (i.e. online calibration). Gas consumption was determined for each bioreactor using gas calibration data. Volumetric consumption rates (mmol/L/day) were calculated by considering exact gas composition, bioreactor medium working volume, feed-and off-gas flow rates, and molar volume of gas. Extracellular products were analysed using filtered broth samples. The culture’s organic acids, alcohols, and fatty acids from 48 and 102 h were quantified by ion-exclusion chromatography using a Vanquish UHPLC system (Thermo Scientific, USA).

### Metagenomic assembly, binning, and annotation

Microbial populations present in wetland study locations and enrichment cultures were characterised through metagenomic binning. The Metaphor pipeline [40] was employed for read quality control, assembly, and binning. Specifically, raw reads from the four metagenome libraries underwent quality control by trimming primers and adapters, removal of artifacts and low-quality reads using fastp [41] with parameters length_required: 50, cut_mean_quality: 30, and extra: --detect_adapter_for_pe. The four metagenomes of each dataset were co-assembled using MEGAHIT v1.2.9 [42] with default settings. Contigs shorter than 1,000 bp were discarded. The assembled contigs were binned with Vamb v4.1.3 [43], MetaBAT v2.12.1 [44], and CONCOCT v1.1.0 [45]. The three binned datasets were subsequently refined using DAS Tool v1.1.6 [46] and dereplicated with dRep v3.4.2 [47] using the parameters -comp 50 -con 10 and default 95% average nucleotide identity. Bin completeness and contamination were assessed using CheckM2 [48]. Quality thresholds for bins were selected based on a previous study [49], retaining only medium (completeness >50%, contamination <10%) and high (completeness >90%, contamination <5%) quality bins for further processing, which were termed metagenome-assembled genomes (MAGs). CoverM v0.6.1 (https://github.com/wwood/CoverM) ‘genome’ was used to calculate the relative abundance of each MAG. MAG taxonomy was determined using the Genome Taxonomy Database Release R214 [50] via GTDB-Tk v2.3.2 [51]. Open reading frames (ORFs) were predicted using Prodigal v2.6.3 [52], then annotated using DIAMOND blastp [53] homology-based searches against a custom database of 51 metabolic marker gene sets [37]. DIAMOND mapping was performed with a query coverage threshold of 80% for all databases, and a percentage identity threshold of 80% (for *psaA*) 75% (for *mcr, hbsT*), 70% (for *isoA, psbA, ygfK, aro, atpA*), 60% (*amoA*, *pmoA*, *mmoA*, *coxL*, *[FeFe]*, *nxrA*, *rbc, nuoFL*) or 50% (all other databases).

### Acetogen metabolic and phylogenetic analyses

Protein sequences of genes involved in the Wood-Ljungdahl Pathway and energy conservation from the genomes of *Acetobacterium woodii, Thermoanaerobacter kivui* [54], and *Sporomusa ovata* [55] were used to create a DIAMOND database [36] (using the parameter “makedb”). This database was used for homology-based identification of protein sequences of genes associated with acetogenesis capabilities using DIAMOND (version 2.0.14) with the parameter “--max-hsps 1 --max-target-seqs 1 --outfmt 6”. A quality control cut-off with a minimum alignment length of 80% and minimum identity threshold of 40% was applied. Data visualization was conducted using custom Python scripts in combination with Adobe Illustrator v29.01. MUSCLE [56] was used to align 416 AcsB protein sequences retrieved between the MAGs and unbinned contigs. A phylogenetic tree was generated using IQ-TREE v2.2.2.6 [57, 58] with 1,000 ultrafast bootstraps [59] and the model LG+F+I+R9. The phylogenetic tree was plotted using iTOL v6 [60].

## Results and Discussion

### Chemosynthetic microbes are abundant in various Blue Carbon ecosystems

Previous observations indicate that the subtropical Australian tidal wetlands under investigation act as a net CO_2_ sink, with fluxes ranging from -0.59 to -0.25 g m^2^ day^-1^ [12]. These fluxes are comparable to those reported for a range of other salt marshes and wetland soils (**Fig. 1a**). The vegetation of Blue Carbon ecosystems is generally efficient at harvesting light for photosynthesis [61], thus strongly limiting light penetration through the canopy and consequently inhibiting CO_2_ uptake through photosynthetic carbon fixation at the soil layer. Therefore, we investigated whether the CO_2_ fluxes we observed in the studied wetland could be mediated by chemosynthetic microbial communities that use reduced chemical compounds as energy sources for carbon fixation and, consequently, sequestration.

**Fig 1.**
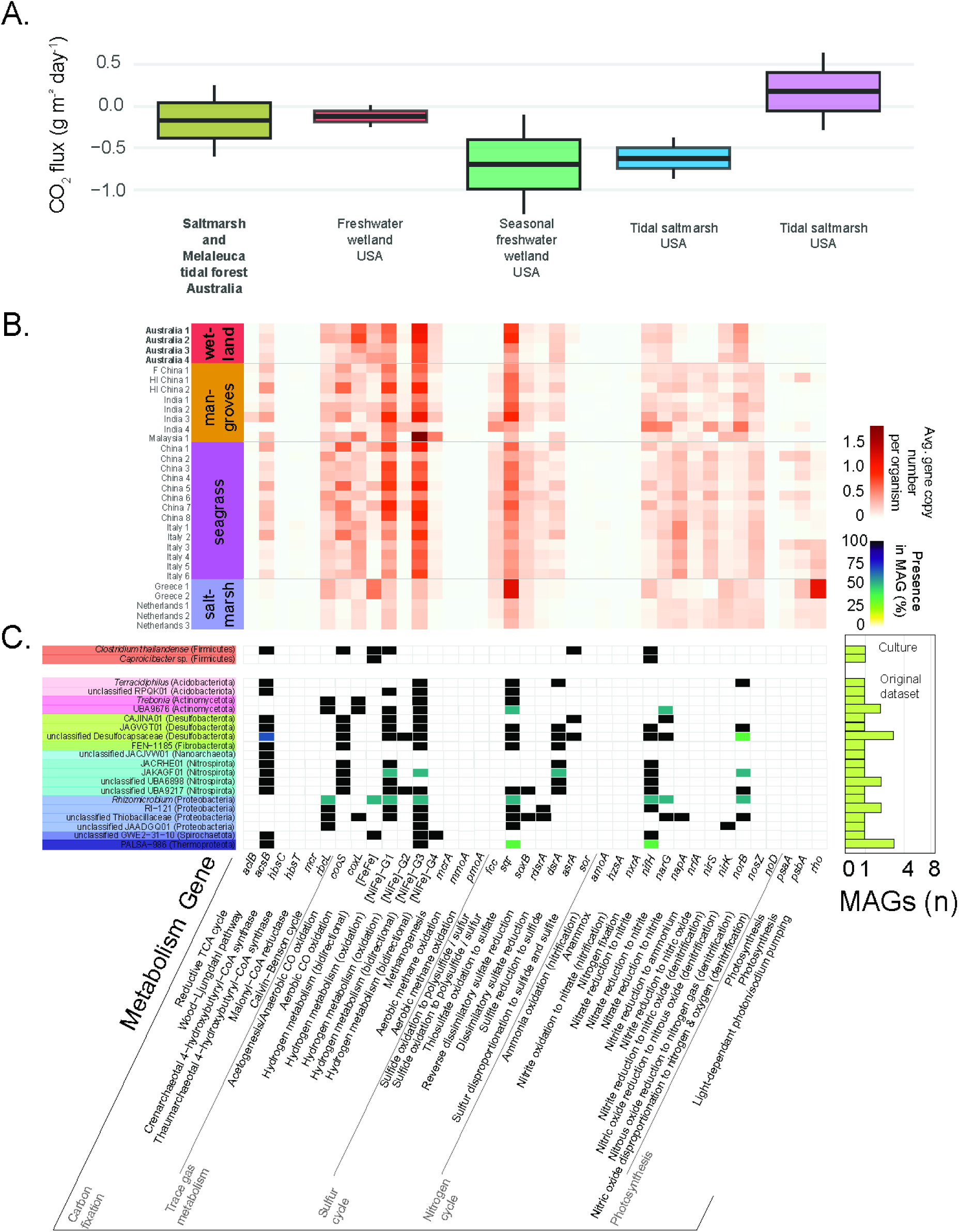
**(A)** CO_2_ fluxes in the studied Blue Carbon ecosystem (bold) compared to other global ecosystems **(B)**. Abundance of metabolic genes from the studied Blue Carbon ecosystem (bold) and various worldwide ecosystems. The heatmap shows the proportion of the community predicted to encode each metabolic marker gene, based on metagenomic sequencing normalized by gene length and the abundance of single-copy marker genes. **(C)**. Presence of metabolic genes in metagenome-assembled genomes (MAGs) from our study site that encode carbon fixation pathways. The top heatmap is for MAGs obtained from the enrichment cultures while the bottom heatmap is for MAGs obtained from the four wetland metagenomes.

First, we sequenced four metagenomes from two locations at the focal site, namely a tidal salt marsh and emerging *Melaleuca* woodland, each at depths of 5 and 7.5 cm **(Fig. S1)**. We conducted a gene-centric analysis to understand which metabolic pathways support energy conservation and carbon fixation in these samples (**Fig. 1b**, **Table S1)**. We found that the capacity for microbial photosynthesis was low (genes encoded by an average of 0.76% of the community), suggesting the salt marsh vegetation and *Melaleuca* trees dominate photosynthesis at these sites. However, there was a very high capacity for carbon fixation through chemosynthetic processes, with a large proportion of the community encoding the marker genes for the Calvin-Benson-Bassham cycle (*rbcL*, av. 21.15% community), Wood-Ljungdahl pathway (*acsB*, 14.1%), reverse tricarboxylic acid cycle (*aclB*, 1.0%), and 3-hydroxypropionate cycle (*mcr*, 0.5%). Consistently, we also observed an extraordinarily high capacity to oxidise inorganic compounds, including hydrogen (group 1 and 2 [NiFe]-hydrogenases, 56% community), sulfide (*sqr*, 64%; *rdsrA*, 4.8%), thiosulfate (8.0%), carbon monoxide (*coxL*, 54%; *cooS*, 23.9%), and iron(II) (*cyc2*; 24.7%). This suggests that a wide portion of the community uses inorganic compounds to derive electrons that sustain respiration and, in some cases, carbon fixation. In line with these sites being variably oxygenated due to tidal flows and soil depth [62], almost all microbes present are predicted to mediate aerobic respiration using both heme-copper and cytochrome *bd* oxidases, but with a strong capacity also to mediate denitrification (*narG*, 20.5%; *napA*, 4.1%), sulfate reduction (*dsrA*, 23.1%; *asrA*, 3.8%), and hydrogenogenic fermentation (group 3b [NiFe]-hydrogenases, 44%; [FeFe]-hydrogenases, 31%) (**Fig. 1b**). Together with the gases CO_2_, H_2_, and CO at these sites, these results suggest there is vast capacity for cycling of nitrogen, sulfur, and carbon compounds, likely mediated by metabolically flexible microorganisms engaging in various biogeochemical interactions.

To generalize these findings, we compared the metagenomes from our study site to those previously sequenced from other Blue Carbon ecosystems, namely salt marshes, seagrass meadows, and mangroves from Europe and Asia [32–35, 63–66] (**Fig. 1b**; **Table S1)**. These sites greatly varied in their capacity for microbial photosynthesis (av. 8.0% community; varying from 0.05% in a Malaysian mangrove [65] to 23.4% in a seagrass meadow of China’s Yellow River [33]), suggesting this is a major process in many Blue Carbon ecosystems though likely less so at our field site. However, the capacity for chemosynthesis was consistently high, with 18.2% encoding the signature gene acetyl-CoA synthase for the WLP (range 0.7% to 45.2%), reverse tricarboxylic acid cycle (av. 2.8%; range 0.5% to 20.1%) and the 3-hydroxypropionate/4-hydroxybutyrate pathway of ammonia-oxidizing archaea (1.5%; range 0% to 4.9%). In certain sites, there was also a high capacity for carbon fixation through the chemosynthetic RuBisCO lineage IE (range 0.0% to 8.5%). The dominant genes related electron donors were predicted to be sulfide (33%), hydrogen (29%), carbon monoxide (26%), thiosulfate (17.8%), iron(II) (5.4%), and ammonia (1.4%). Whereas the dominant genes related to electron acceptors were predicted to be oxygen (100%), nitrate (21%), organohalides (13.7%) and sulfate (7%) with hydrogen as a key fermentative sink (66.2%) **(Table S1, Fig. 1b**).

These findings suggest that the high genetic capacity for biogeochemical cycling and chemosynthetic carbon fixation at our field site are extendable to other Blue Carbon ecosystems, although the key processes and mediators vary. There is likely much cryptic elemental cycling in these ecosystems, for example, through the coupling of fermentation with hydrogen oxidation, sulfate reduction with sulfide oxidation, and organic carbon oxidation with carbon fixation, in addition to the net consumption of atmospheric carbon dioxide and likely other trace gases. These findings indicate Blue Carbon ecosystems are broadly hotspots for biogeochemical cycling marked by a remarkable co-existence of photosynthetic and chemosynthetic processes, as well as aerobic respiratory, anaerobic respiratory, and fermentative metabolisms.

### Phylogenetically and metabolically diverse acetogens inhabit tidal wetlands

To better understand the mediators of chemosynthesis in Blue Carbon ecosystems, we reconstructed 29 archaeal and 88 bacterial metagenome-assembled genomes from the Australian tidal wetland, which collectively mapped on average to 17% of metagenomic reads **(Table S2)**. In line with the predictions based on metabolic short reads (**Fig. 1b**), there was a vast capacity for aerobic respiration, denitrification, sulfate reduction, lithotrophy, and fermentation (**Fig. 1c**). Most mediators of these processes were from uncultivated genera and phyla, including numerous novel archaea. In line with inhabiting a variably oxygenated environment, almost all bacterial MAGs are predicted to be capable of aerobic respiration, including all MAGs predicted to mediate dissimilatory sulfate reduction and 13 of the 17 MAGs that encode the WLP; this suggests bacteria are capable of aerobic growth and a high aerotolerance of obligate anaerobes (**Fig. 1c**). Overall, the bacteria inhabiting these environments are metabolically flexible, capable of using a wide range of electron donors, carbon sources, and electron acceptors. Such observations are reminiscent of previous findings that metabolic flexibility enables microbes to become dominant in tidally disturbed sandy soils [67, 68]. Nevertheless, the investigated site could also provide diverse spatiotemporal niches that enrich for obligate anaerobes.

Multiple MAGs were predicted to be capable of chemosynthesis, though none encoded photosynthesis genes. Seven bacterial MAGs encoded RuBisCO, namely from the lineages Streptosporangiales, Burkholderiales, Micropepsales, and order JAADGQ01; these are each predicted to be aerobic lithoautotrophs capable of using multiple inorganic energy sources, namely sulfide (6 MAGs), hydrogen (5 MAGs), and carbon monoxide (4 MAGs) (**Fig. 1c**). In addition, 18 MAGs from eight phyla encoded the acetyl-CoA synthase for the WLP (**Fig. 1c**, **Fig. 3a**), often in concert with carbon monoxide dehydrogenases (10 MAGs) and hydrogenases (9 MAGs). We reconstructed the metabolic capabilities of these MAGs and confirmed they each encoded a largely complete WLP (**Fig. 3b**). No MAGs capable of other carbon fixation pathways were constructed.

### Enrichment cultures grown on carbon oxides and hydrogen suggest Clostridial species encode and mediate hydrogenotrophic acetogenesis

We subsequently validated that acetogens are present and active in Blue Carbon ecosystems by culturing them in chemically defined media with carbon oxides and H_2_ as the sole carbon and energy source. We exposed soil samples to three distinct gas conditions (CO_2_/H_2_, CO/CO_2_/H_2_, and CO under anoxic conditions), and only cultures grown on CO_2_/H_2_ showed notable growth and decrease in gas pressure (**Fig. 3a, 3b**).

Following anaerobic enrichment by four iterative in liquid **Fig. 3c, 3d**) and solid culture, two putatively acetogenic cultures were obtained (hereafter named A and B) that grew to high optical densities on CO_2_/H_2_ (OD_max_ = 0.39 and 0.37, respectively) and consumed much of the gas (33% and 43% of the initial pressure).

Metagenomic sequencing confirmed these enrichment cultures predominantly comprised the bacteria *Clostridium thailandense* (78% and 26% reads) and *Caproicibacter* (comprising 4.2% and 65% reads) (**Fig. 3f**). Though both bacteria encoded [FeFe]-hydrogenases capable of H_2_ uptake, only the *C. thailandense* isolate encoded carbon fixation/acetogenesis genes (**Fig. 3e**), as supported by phylogenetic analysis (**Fig. 2a**). Hence, this bacterium likely accounts for the observed conversion of H_2_ and CO_2_ to acetate. The same species was previously isolated from peatland soil, where it was demonstrated to mediate hydrogenotrophic acetogenesis [69]. Read mapping confirmed that this bacterium occurred in all four tidal wetland samples subject to metagenomic sequencing and, hence, likely contributes to chemosynthesis at these sites **(Table S3).**

**Fig 2.**
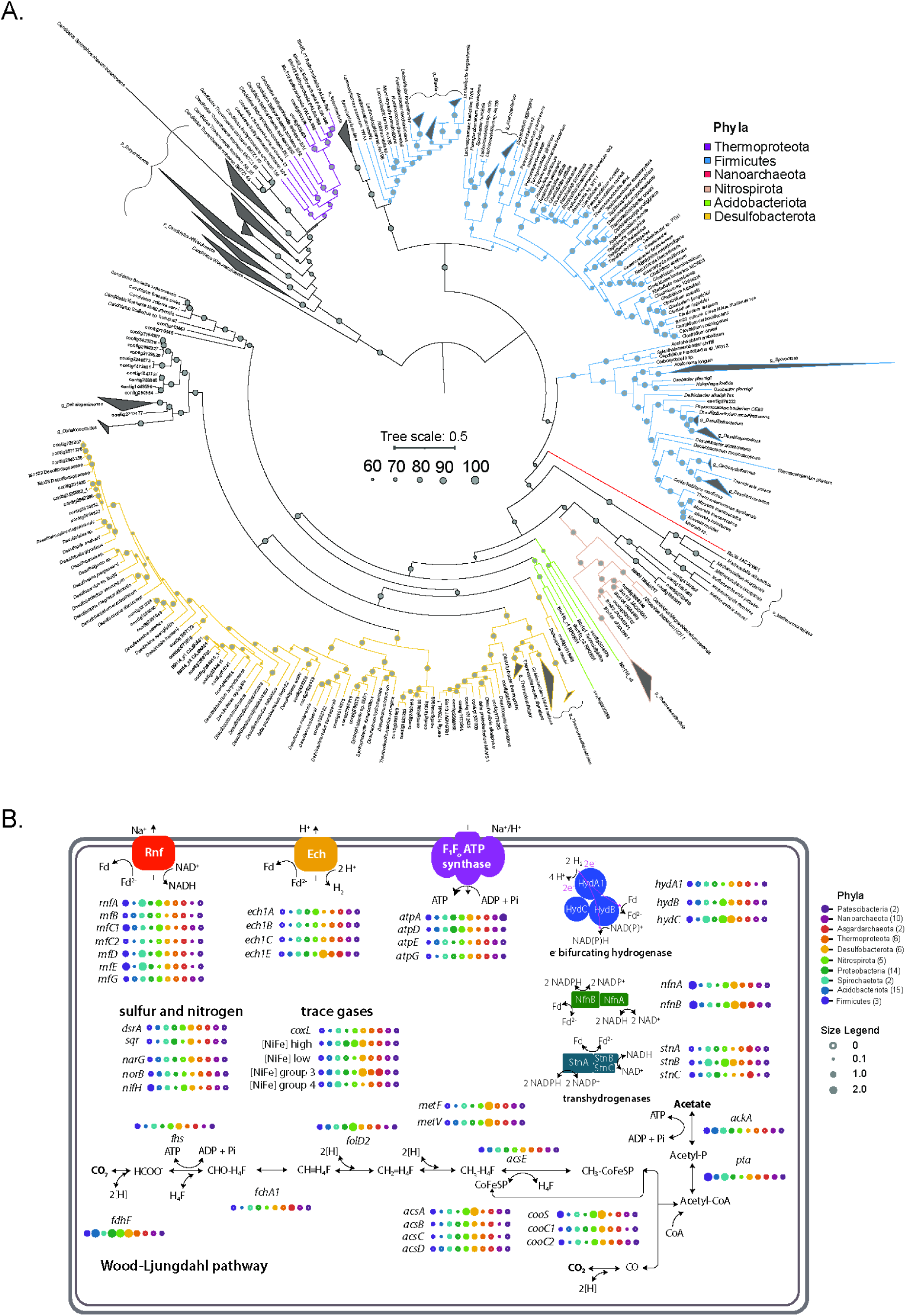
**(A)** Maximum-likelihood phylogenetic tree of the acetyl-CoA synthase (AcsB) protein. Shown are sequences from metagenome-assembled genomes (MAGs) and contigs (bold). Amino acid sequences of MAGs and contigs were aligned with reference sequences, and the tree was bootstrapped with 1000 replicates. **(B)** Metabolic diagram illustrating chemosynthetic potential in ten phyla that encode genes of the Wood-Ljungdahl pathway. The number of genomes affiliated with the respective phylum is indicated in brackets. The abundance of each gene was calculated as the phylum-level mean copy number per organism (circles), differentiated by colour.

The metabolic capabilities of *C. thailandense* were predicted by annotating a 99.99% complete and 5% contaminated MAG (**Fig. 3e**). The bacterium encoded a near-complete WLP, including both the carbonyl branch and the methyl branch to generate acetyl-CoA, though lacked a canonical methylene-tetrahydrofolate reductase (**Fig. 3e**). Its energy conservation mechanism resembles the model acetogen *Clostridium ljungdahlii*, relying on the membrane-bound Rnf complex that couples electron transfer from reduced ferredoxin to NAD^+^ with translocation of protons across the cytoplasmic membrane to generate a proton-motive force that drives ATP synthesis. This bacterium is also predicted to use an electron-bifurcating [FeFe]-hydrogenase (HydABC) to generate reductant (NAD[P]H and reduced ferredoxin) from hydrogen, as well as the electron-bifurcating enzyme Nfn to catalyse the interconversion of ferredoxin, NADH and NADPH. Interestingly, this bacterium also encodes a respiratory group 1a [NiFe]-hydrogenase and anaerobic sulfite reductase (Asr), suggesting it potentially conserves energy by hydrogenotrophic sulfate reduction simultaneously or with acetogenesis. *C. thailandense* also encodes a nitrogenase, which likely enabled it to grow in the absence of a fixed nitrogen source in the enrichment cultures (**Fig. 3e**). Altogether, these findings suggest that acetogenesis can play a role in micro-aerobic environments, contributing to diverse consortia and carbon sequestration into Blue Carbon ecosystems.

**Fig 3.**
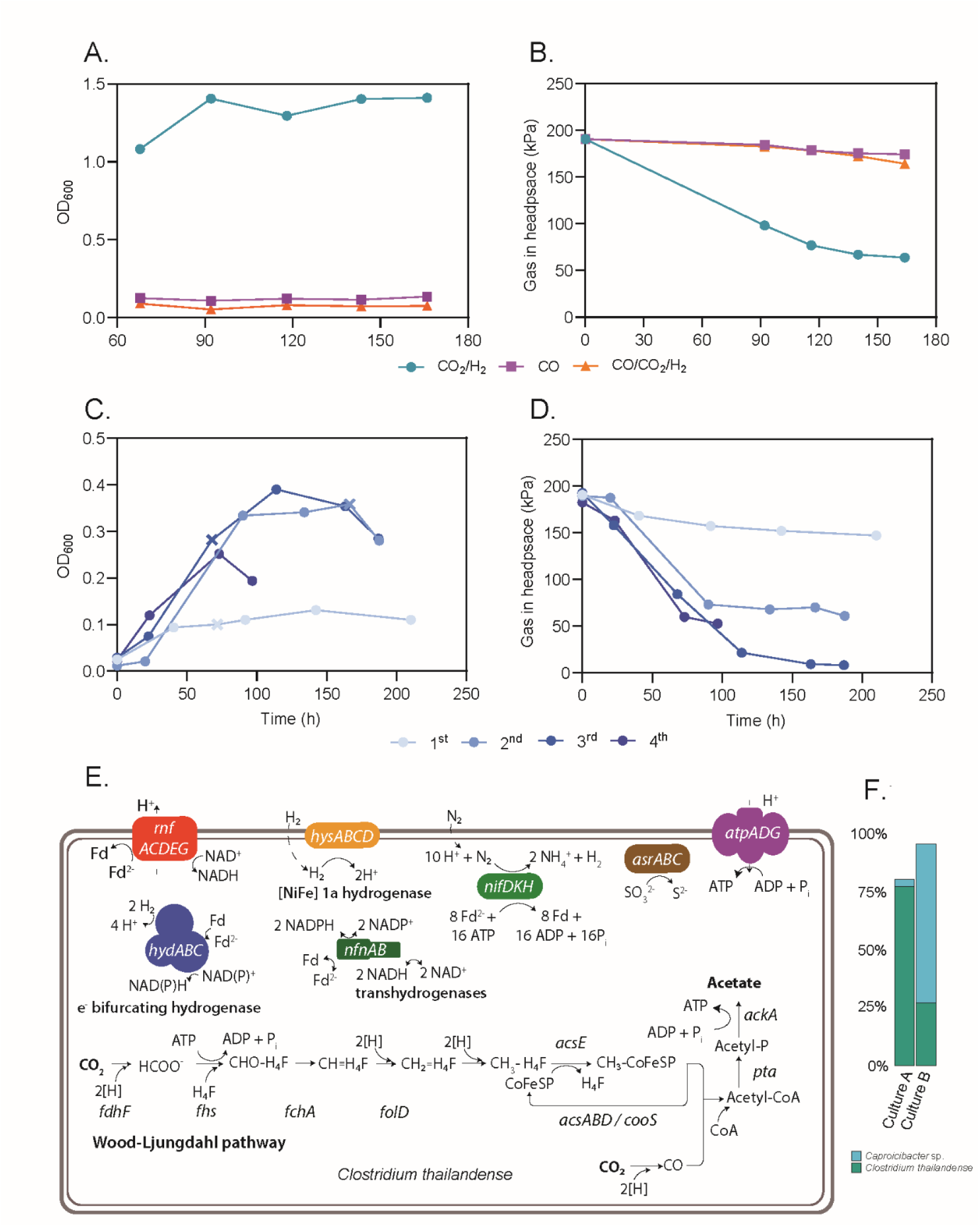
**(A-B)** Bacterial growth (A) and gas consumption (B) of enrichment cultures from the tidal wetland soils. Cultures were exposed to three different gas mixtures: CO_2_/H_2_, CO, and CO/CO_2_/H_2_. **(C-D)** Bacterial growth (C) and headspace gas pressure (D) in four iterative CO_2_/H_2_ subcultures of the original CO_2_/H_2_ culture. X symbols represent the sample used as inoculum for consecutive subcultures. **(E)** Metabolic reconstruction of *Clostridium thailandense* illustrating the presence of genes involved in acetogenesis. Methylene-tetrahydrofolate reductase completes the reaction that is not labelled, i.e., between *folD* and *acsE*. Abbreviations: *cooS* – carbon monoxide dehydrogenase can be in complex with, *acsABD* -acetyl-CoA synthase; *fdhF*– formate dehydrogenase can be in complex with, *hydABC* – electron bifurcating [FeFe]-hydrogenase; *fhs* – formate:THF ligase; *fchA* – 5,10-methenyl-THF 5-hydrolase; *folD* – methylene-THF dehydrogenase; *pta* – phosphotransacetylase; *ackA* – acetate kinase; *rnfACDEG –* Rnf complex*; NfnAB* – Nfn transhydrogenase complex; hysABCD – [NiFe] 1a hydrogenase; nifDKH – nitrogenase complex; asrABC – sulfite reduction complex; atpADG – ATPase synthase. **(F)** Proportion of species present in enriched cultures A and B grown on CO_2_/H_2_.

### Enriched cultures rapidly convert carbon dioxide to fatty acids and ethanol

To demonstrate the successful isolation of species that can grow autotrophically, we grew enrichment cultures A and B in CO_2_/H_2_ batch bioreactors (**Fig. 4a, 4b**). Both cultures demonstrated rapid growth in line with gas uptake. Compared to small-scale cultures in serum bottles, bioreactors enabled higher biomass concentrations (OD_max_ = 0.75 and 0.62) attributed to continuous gas substrate supply and controlled culture conditions. Both cultures started consuming CO_2_ and H_2_ after ∼24h from inoculation in parallel to exponential growth. During this growth phase, cultures A and B exhibited growth rates of 0.039 and 0.029 h^-1^. Maximum gas consumption rates were determined towards mid-exponential growth, when H_2_ became the preferred substrate, likely related to the activity of hydrogenases present in both *C. thailandense* and *Caprocibacter* sp. The enriched culture A (mainly comprised of *C. thailandense*) showed improved culture performance, reaching higher biomass concentrations and consistency in gas consumption aligned with growth. Strikingly, culture B showed fluctuating patterns in gas consumption. The reasons for these oscillations are unknown but might be related to CO_2_/H_2_ consumption and production mechanisms resulting from the consortium of *Caproicibacter* and *C. thailandense* ratio within this culture, therefore resulting in impaired growth patterns compared to culture A.

**Fig 4.**
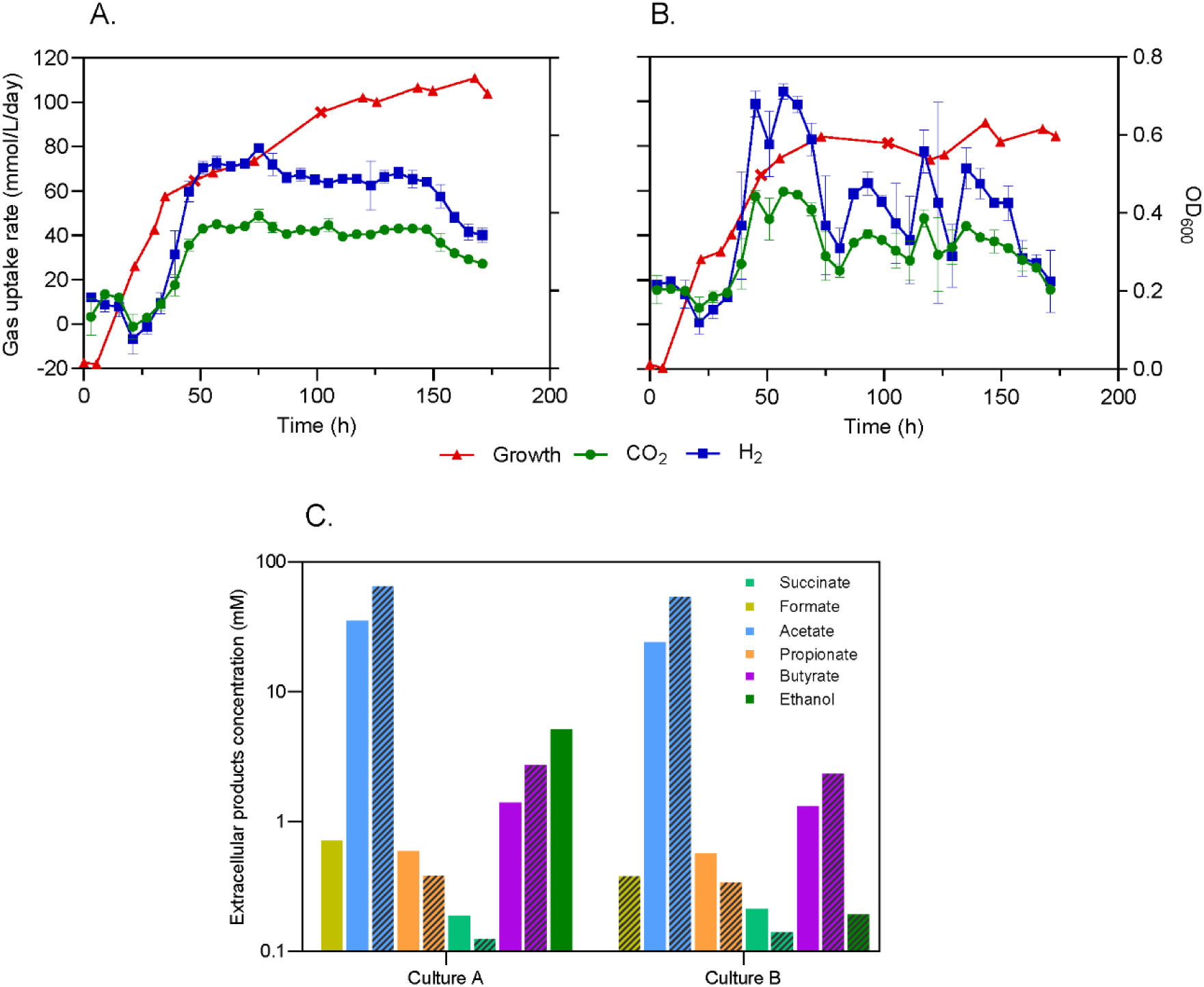
**(A-B)** Bacterial growth (OD_600_) and volumetric CO_2_/H_2_ gas consumption from cultures A (A) and B (B) in batch bioreactor fermentation. Extracellular product sample time points are illustrated with X-symbols. Volumetric gas uptake rates were determined as a mean ± standard deviation (error bars) over a 5 h period. **(C)** Extracellular product quantification during exponential (48 h, solid) and stationary (102 h, striped) phases.

Production of known microbial anaerobic products [70], such as acetate, propionate, succinate and butyrate, was observed in both bioreactor cultures (**Fig. 4c**). Significant concentrations of acetate, the main metabolic product of the WLP, were quantified in both cultures (64 and 53 mM in culture A and B). Additionally, culture A produced other common acetogenic products, such as formate during exponential growth and ethanol during stationary phase. In culture B, these compounds were quantified during the stationary phase only at lower concentrations, likely suggesting production from *C. thailandense*. Butyrate titers increased similarly for stationary phase in both cultures, indicating improved medium-chain carboxylate elongation as biomass and availability of ethanol and acetate increased. In agreement with our findings, butyrate biosynthesis via butyryl-CoA has been extensively reported in *Clostridium* [71–74] and *Caproicibacter* [75, 76] species. Decreased propionate and succinate were determined at the later stage of culture, indicating their possible use as intermediates for biomass generation and further chain elongation. Similar to our findings, propionate production was demonstrated from an enriched homoacetogen consortium (from peatland soil) when grown with CO_2_/H_2_ [77].

Beyond demonstrating the potential of acetogens in carbon sequestration within wetlands, the enrichment of microbial cultures revealed the importance of microbial interactions in blue carbon ecosystems. Furthermore, bioreactor-based gas fermentation provided valuable insights into the metabolic traits of these microorganisms. Such knowledge is essential for the isolation of further optimization of robust CO_2_-fixing microbial strains, enabling their application in gas fermentation for the sustainable production of high-value products

## Conclusions

Overall, our study emphasizes the role of Blue Carbon ecosystems in carbon capture and storage and highlights the contribution of microbial communities in carbon sequestration within these ecosystems. Tidal wetlands harbor highly diverse archaeal and bacterial communities with flexible metabolic capabilities to use a wide range of carbon sources, electron donors, and electron acceptors. Our results indicated the presence of microbial communities with the ability to fix CO_2_ through chemosynthetic metabolic pathways, including both aerobic and anaerobic lithoautotrophs encoding the CBB cycle and WLP, respectively. In addition to current knowledge on acetogenic lineages, findings from our metabolic reconstructions expand the knowledge on acetogenic bacterial capabilities and suggest novel diversity of archaea able to mediate acetogenesis in Blue Carbon ecosystems.

Incorporating the role of chemosynthetic microorganisms in carbon sequestration and their potential applications in gas fermentation can enhance Blue Carbon initiatives, accounting for the broader range of ecological processes that contribute to carbon storage in systems [12, 13]. This approach not only improves the accuracy of carbon sequestration assessments, but also uncovers novel strategies for boosting the efficacy of Blue Carbon projects and supporting sustainable environmental management. Future research should examine how tidal wetlands, beyond sequestering carbon in soil and woody biomass, may also directly consume CO_2_. This insight could inspire innovative gas fermentation applications replicating natural processes in tidal wetlands.

## Study funding

This research was funded by the Australian Government through the Australian Research Council Centre of Excellence in Synthetic Biology (CE 200100029) and the Woodside-Monash Energy Partnership.

## Conflict of interest

The authors declare no conflict of interest.

## Data availability statement

The metagenomic reads and MAGs generated during and/or analysed in the current study are available in the NCBI Bioproject: PRJNA1184823 and supplementary files.

## Acknowledgments

We thank Julieta Gamboa and Damien Maher from the Australian Rivers Institute for the field work and providing of the soil samples from tidal wetlands. We acknowledge Queensland Metabolomics and Proteomics (Q-MAP) for providing analytical results and IDEA Bio for their support with bioreactor design. Q-MAP and IDEA Bio are supported by Bioplatforms Australia (BPA), a National Collaborative Research Infrastructure Strategy (NCRIS) funded facility. We also acknowledge the support from the Australian Centre for Ecogenomics (ACE) and the Ramaciotti Centre for Genomics for providing metagenomic sequencing data.

## Author contributions

KR: Conceptualization, data curation, formal analysis, investigation, methodology, visualization, writing – original draft, writing – review and editing. FR: Conceptualization, data curation, formal analysis, investigation, methodology, visualization, writing – original draft, writing – review and editing. GN: data curation, formal analysis, investigation, methodology, visualization, writing – review and editing. NI: Methodology, writing – review and editing. RP: Methodology, writing – review and editing. RAGG: Methodology, writing – review and editing. JH: Methodology, writing – review and editing. MFA: Conceptualization, methodology, writing – original draft, writing -review and editing. CG: Conceptualization, data curation, formal analysis, investigation, methodology, visualization, writing – original draft, writing – review and editing. EM: Conceptualization, funding acquisition, investigation, supervision, writing – original draft, writing – review and editing.

## Notes

### Competing Interest Statement

The authors have declared no competing interest.

