## Supplementary figures and images for "Abundant and active acetogens enhance the carbon dioxide sink of Blue Carbon ecosystems"

### Fig. S1

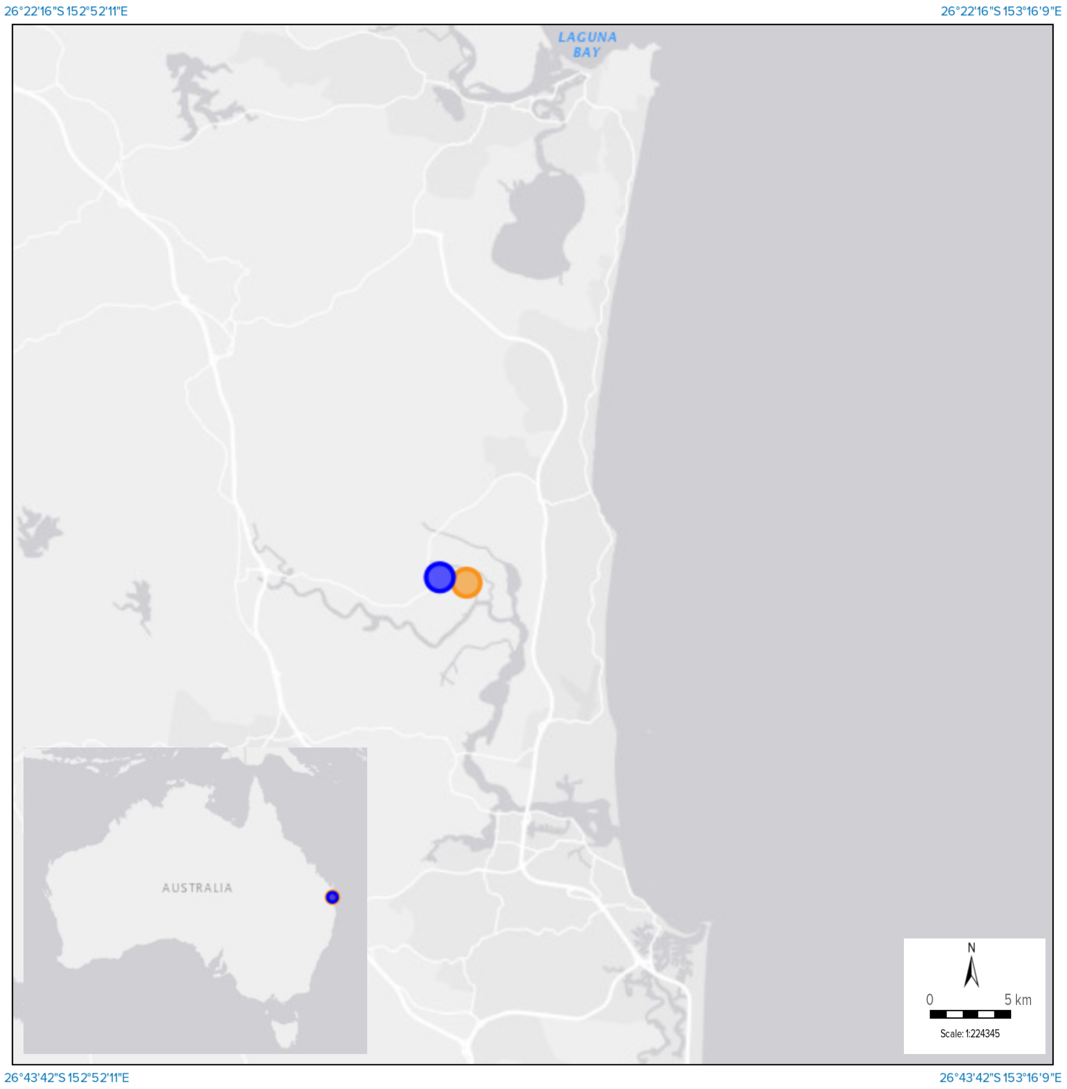

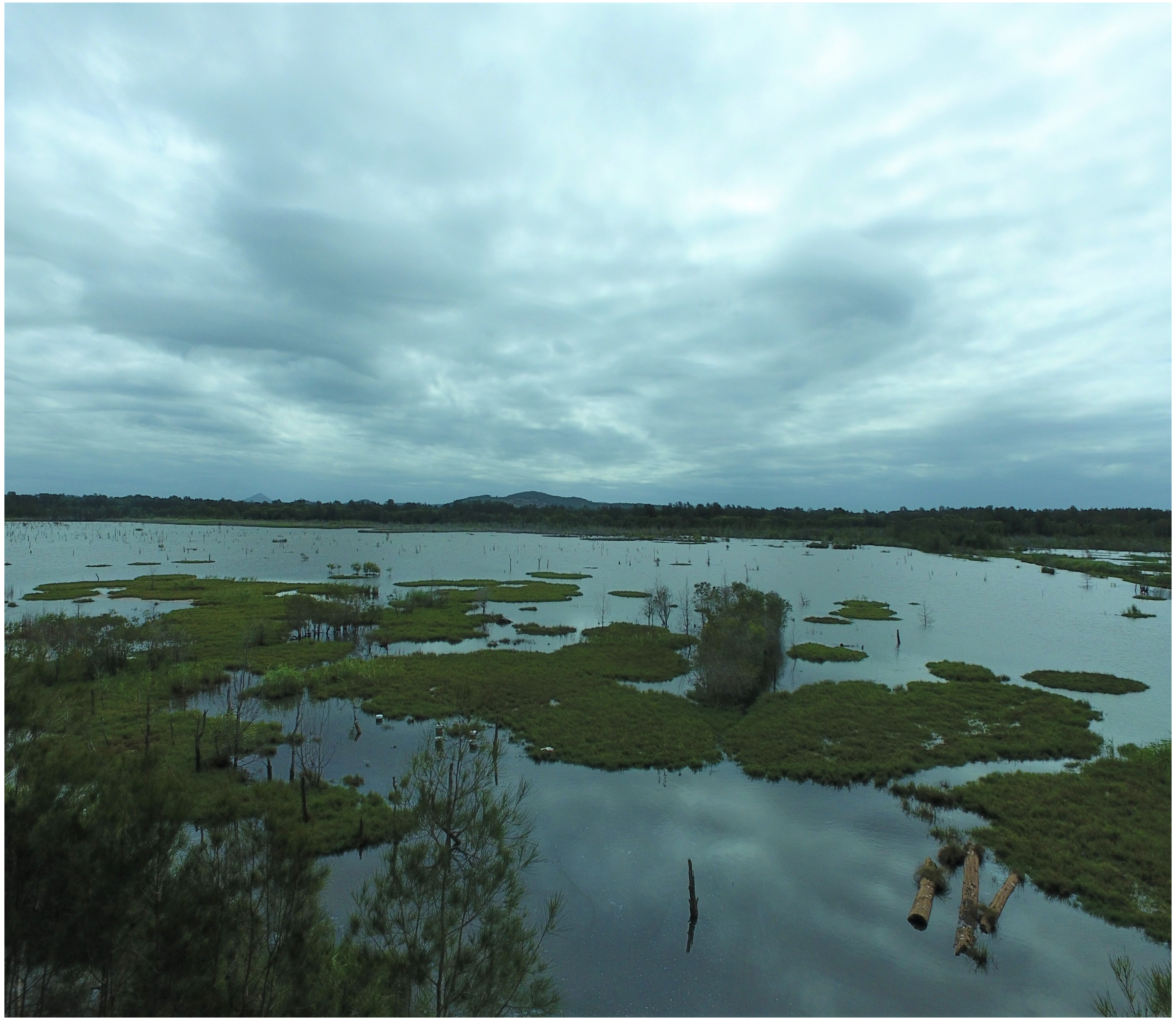
